# Inhibiting glucocorticoid receptors enhances adult spinal cord neural stem cell activity and improves outcomes in spinal cord injury

**DOI:** 10.1101/2025.05.06.652540

**Authors:** Xuefeng Zhang, Siyuan Zhou, Shengyu Tang, Xiaotong Hou, Yonghui Cai, Changlong Hu

**Author notes:** **Corresponding author:** Changlong Hu.

## Abstract

The internal microenvironment plays a critical role in the proliferation and differentiation of endogenous neural stem/progenitor cells (NSPCs). A big change in the spinal cord injury (SCI) microenvironment is the elevated level of glucocorticoids. In this study, we examined the impact of glucocorticoids on endogenous NSPCs in the adult mouse spinal cord. Our findings reveal that adult spinal cord NSPCs express glucocorticoid receptors, but not mineralocorticoid receptors. Glucocorticoids were found to significantly inhibit the proliferation and neurosphere formation of NSPCs via activation of glucocorticoid receptors, and they also impaired their differentiation. Importantly, the glucocorticoid receptor inhibitor CORT125281 was shown to enhance motor function in a traumatic SCI model in mice. Treatment with CORT125281 increased the number of NSPCs at the injury site in vivo. Flow cytometry and RNA sequencing analyses indicated that glucocorticoids induce NSPC arrest in the G1/G0 phase through the p53 signaling pathway. Glucocorticoids increased the expression of cell-cycle regulatory genes p15, p16, p18, and p27 in adult spinal cord NSPCs. In summary, our data suggest that glucocorticoids elevation following SCI suppresses the proliferation of endogenous NSPCs via glucocorticoid receptor activation. Targeting glucocorticoid receptors with specific inhibitors may represent a novel therapeutic strategy to promote recovery after spinal cord injury.

## Introduction

Spinal cord injury (SCI) is a complex and severe neurological condition, typically resulting from traumatic events such as traffic accidents and falls. This condition often leads to debilitating neurological deficits, including loss of sensation, bowel and bladder dysfunction, paraplegia, and tetraplegia ^1^. Despite extensive research efforts, effective neurorestorative treatments for SCI remain elusive, with current interventions focusing primarily on symptomatic management and physical rehabilitation, offering limited functional recovery. Stem cell transplantation has been extensively explored as a potential therapeutic approach for SCI. Among the various types of stem cells studied, mesenchymal stem cells have emerged as particularly promising candidates ^2,3^. A recent Phase I clinical trial demonstrated that intrathecal injection of adipose-derived mesenchymal stem cells in ten patients with traumatic SCI resulted in no serious adverse events, with seven of the ten patients showing an improvement in their American Spinal Injury Association Impairment Scale grade compared to their pre-injection status ^4^.

In addition to stem cell transplantation, there is growing evidence supporting the activation of endogenous neural stem/progenitor cells (NSPCs) as a promising strategy for SCI repair ^5,6^. Adult endogenous spinal cord NSPCs, derived from the ependymal layer of the central canal, play a crucial role in promoting functional recovery post-SCI ^7^. Various approaches have been explored to stimulate endogenous NSPCs, including transplanting biomaterials that replicate spinal cord mechanics and using functional biomaterials for sustained drug release, among others ^6^. Regardless of whether exogenous stem cells or endogenous NSPCs are chosen as the repair strategy, both approaches must contend with the significant changes in the microenvironment following SCI.

The spinal cord’s microenvironment, composed of various cells, extracellular matrix (ECM), and signaling molecules, undergoes dramatic changes following injury, which can either promote or hinder the recovery process ^8^. Understanding these microenvironmental changes after SCI is crucial for developing therapeutic strategies that can modulate the post-injury environment to promote regeneration and improve functional outcomes. Previous study showed that glucocorticoid levels were significantly increased in both murine experimental SCI and human SCI ^9^. Glucocorticoids have been shown to inhibit the proliferation of both mouse and rat embryonic NSPCs ^10,11^. These studies suggest that glucocorticoids may also inhibit adult endogenous spinal cord NSPCs and impair SCI repair.

In present study, we investigated the impact of glucocorticoids on endogenous NSPCs in the adult mouse spinal cord. Our data showed that glucocorticoids elevation following SCI suppresses the proliferation of endogenous NSPCs via glucocorticoid receptor activation, and that the glucocorticoid receptor inhibitor CORT125281 enhanced motor function in a mice model of traumatic SCI. Targeting glucocorticoid receptors with specific inhibitors may represent a novel therapeutic strategy to promote recovery after spinal cord injury.

## Results

### Glucocorticoids suppress the proliferation of adult mouse spinal cord NSPCs

To investigate the effect of glucocorticoids on the proliferation of spinal cord NSPCs, we employed a neurosphere assay. Since cortisol is the predominant glucocorticoid in humans and corticosterone in rodents, we assessed the effects of both. Cultured NSPCs were seeded in 96- well plates and treated with proliferation medium containing varying concentrations of cortisol or corticosterone (0.1–50 μM). The number of neurospheres was counted seven days after plating. Both glucocorticoids significantly reduced neurosphere formation in a concentration-dependent manner (Fig. 1A-D). To confirm this inhibitory effect, CCK-8 assays revealed that both cortisol and corticosterone decreased NSPC viability (Fig. 1E and F). Furthermore, EdU incorporation assays showed a significant reduction in EdU-positive NSPCs upon treatment with 1 μM cortisol (Fig. 1G and H). Since 1 μM is a pathologically relevant concentration in both spinal cord injury (SCI) patients and rodent models^9^, this concentration was selected for subsequent experiments.

**Fig. 1.**
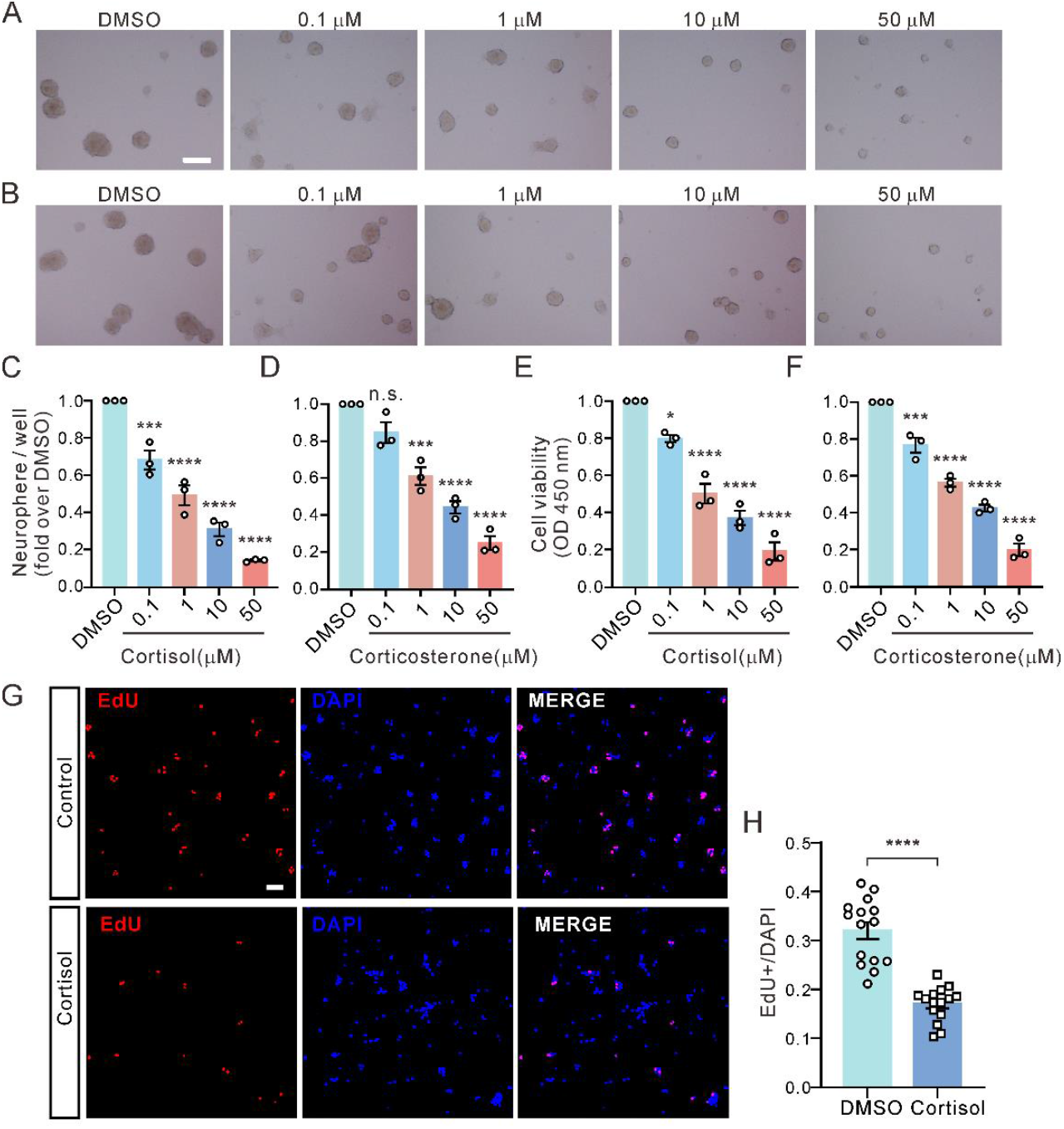
Glucocorticoids suppress the proliferation of adult mouse spinal cord NSPCs. **(A)** Representative images showing the effect of cortisol (1–50 μM) on NSPC neurosphere formation. **(B)** Representative images showing the effect of corticosterone (1–50 μM) on neurosphere formation. Scale bar: 100 μm. **(C)** Statistical analysis of neurosphere formation from adult spinal cord NSPCs with/without various concentrations of cortisol treatment, performed using one-way ANOVA with Bonferroni’s post hoc test (n = 3, F (4, 10) = 81.36, p < 0.0001). ***p = 0.0005; ****p < 0.0001. **(D)** Statistical analysis of neurosphere formation from adult spinal cord NSPCs with/without various concentrations of corticosterone treatment using one-way ANOVA with Bonferroni’s post hoc test (n = 3, F (4, 10) = 57.48, p < 0.0001). n.s., not significant (p = 0.0858); ***p = 0.0002; ****p < 0.0001. **(E)** Statistical analysis of the effect of cortisol (1–50 μM) on the viability of spinal cord NSPCs, performed using one-way ANOVA with Bonferroni’s post hoc test (n = 3, F (4, 10) = 76.29, p < 0.0001). *p = 0.0135; ****p < 0.0001. **(F)** Statistical analysis of the effect of corticosterone (1–50 μM) on the proliferation of spinal cord NSPCs using one-way ANOVA with Bonferroni’s post hoc test (n = 3, F (4, 10) = 136.5, p < 0.0001). ***p = 0.0004; ****p < 0.0001. **(G)** Representative figures showing the effect of 1 μM cortisol on EdU incorporation. Scale bar: 50 μm. **(H)** Statistical analysis of EdU-positive cells from adult spinal cord NSPCs with/without treatment with 1 μM cortisol using a two-tailed unpaired t-test (n = 15, t (28) = 7.687, ****P < 0.0001).

### Glucocorticoid receptors mediate the inhibitory effect of glucocorticoids on adult NSPC proliferation

Glucocorticoid effects can be mediated through glucocorticoid or mineralocorticoid receptors. Previous studies have shown that embryonic neural stem cells express both receptors^12^. Immunofluorescence analysis revealed that cultured adult spinal cord NSPCs express glucocorticoid receptors but do not show detectable mineralocorticoid receptor expression (Fig. 2A). To confirm the specificity of the mineralocorticoid receptor antibody, HeLa cells, which express mineralocorticoid receptors ^13^, were used as a positive control (Fig. 2B). Adult endogenous spinal cord NSPCs originate from the ependymal layer of the central canal of the spinal cord ^7,14^. Further analysis showed that glucocorticoid receptors are predominantly localized in the ependymal layer of the central canal in the adult mouse spinal cord, while mineralocorticoid receptors are absent (Fig. 2C).

**Fig. 2.**
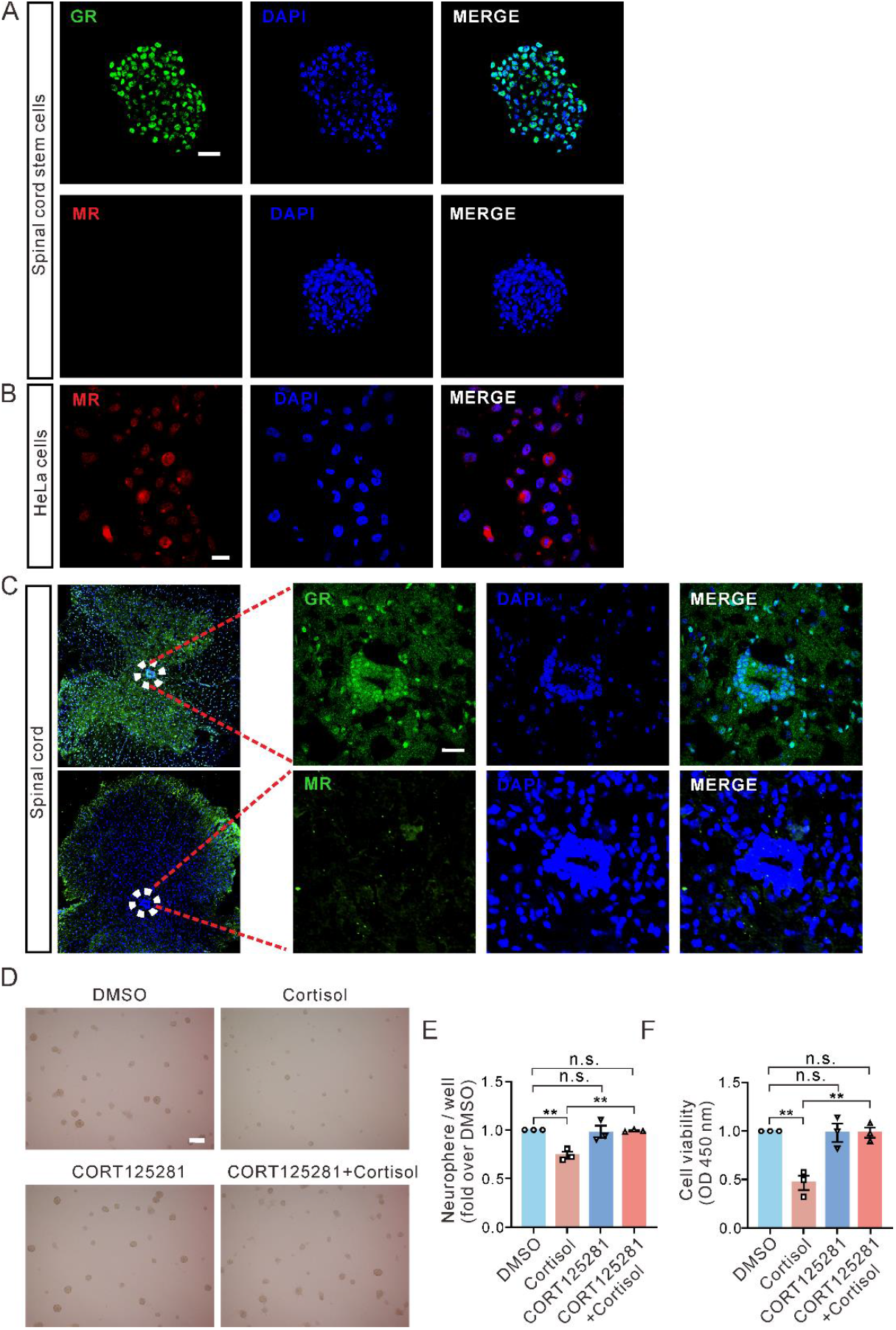
Glucocorticoid receptors mediate the inhibitory effect of glucocorticoids on NSPC proliferation. **(A)** Representative immunofluorescence images showing that spinal cord NSPC neurospheres express glucocorticoid receptors (GRs) but not mineralocorticoid receptors (MRs). **(B)** Representative immunofluorescence images showing the expression of MRs in HeLa cells. **(C)** Representative immunofluorescence images showing that the central canal region of mouse spinal cord tissue expresses GRs but not MRs. **(D)** Representative images of neurosphere formation from adult spinal cord NSPCs pre-incubated with 500 nM CORT125281 for 30 minutes, followed by incubation with or without 1 μM cortisol. **(E)** Statistics for the number of neurospheres from D using one-way ANOVA with Bonferroni’s post hoc test (n = 3, F (3, 8) = 11.49, p = 0.0029). Comparisons: DMSO vs. cortisol: **p = 0.0054, DMSO vs. CORT125281 + cortisol: p > 0.9999, DMSO vs. CORT125281: p > 0.9999, cortisol vs CORT125281 + cortisol: **p = 0.0093, n.s., not significant. **(F)** Statistical analysis of the effect of CORT125281 on the cortisol-induced inhibition of spinal cord NSPC viability, analyzed using one-way ANOVA with Bonferroni’s post hoc test (n = 3, F (3, 8) = 15.47, p = 0.0011). Comparisons: DMSO vs. cortisol: **p = 0.0026, DMSO vs. CORT125281 + cortisol: p > 0.9999, DMSO vs. CORT125281: p > 0.9999, cortisol vs CORT125281 + cortisol: **p = 0.0035; n.s., not significant. Scale bars: 25 μm (A, B, C), 200 μm (D).

To investigate whether the glucocorticoid receptor mediates the effects of glucocorticoids, we used the specific receptor blocker CORT125281^15^. Both the neurosphere assay and CCK-8 experiments demonstrated that pre-incubation with the glucocorticoid receptor-specific antagonist, CORT125281(500 nM) for 30 minutes, abolished the inhibitory effects of cortisol on NSPC proliferation (Fig. 2D-F). These results indicate that glucocorticoids inhibit NSPC proliferation through the glucocorticoid receptor. We next investigated the effects of glucocorticoids on NSPC differentiation.

### Glucocorticoids impair the differentiation of adult spinal cord NSPCs

To assess the role of glucocorticoids in NSPC differentiation, cells were cultured in differentiation medium with or without 1 μM cortisol for seven days. Immunofluorescence analysis showed that cortisol inhibited NSPC differentiation into glial cells, an effect that was partially reversed by the glucocorticoid receptor antagonist CORT125281 (Fig. 3A-B). Cortisol also suppressed neuronal differentiation, although CORT125281 did not affect this inhibition (Fig. 3C-D). Furthermore, glucocorticoids influenced the morphology of neurons. Cortisol not only reduced the number of neurites, an effect that could not be blocked by CORT125281, but also significantly decreased neurite length. This reduction in neurite length was partially reversed by CORT125281 (Fig. 3E-H).

**Fig. 3.**
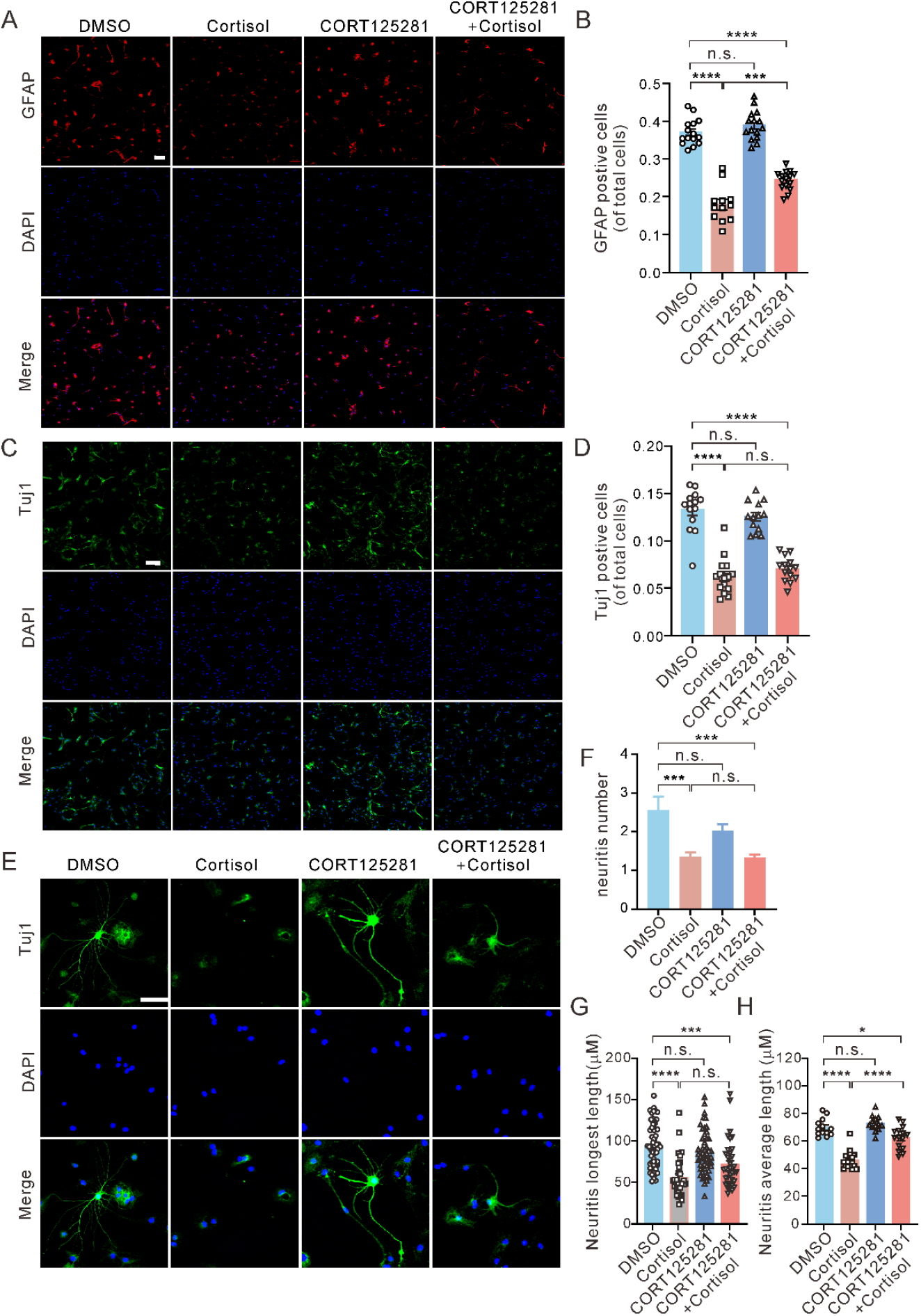
Glucocorticoids impair the differentiation of adult spinal cord NSPCs. **(A)** Representative immunofluorescence images showing the effect of 1 μM cortisol in the presence or absence of CORT125281 on the differentiation of NSPCs. **(B)** Statistical analysis of the number of GFAP (glial cell marker protein)-positive cells from panel A using one-way ANOVA with Bonferroni’s post hoc test (n = 13-15, F (3, 53) = 100.3, p < 0.0001). Comparisons: DMSO vs. cortisol: ****p < 0.0001; DMSO vs. CORT125281 + cortisol: ****p < 0.0001; DMSO vs. CORT125281: p > 0.9999, n.s., not significant; cortisol vs. CORT125281 + cortisol: ***p = 0.0002. **(C)** Representative immunofluorescence images showing the effect of 1 μM cortisol in the presence or absence of CORT125281 on the differentiation of NSPCs. **(D)** Statistics for the number of Tuj1(neural marker protein)-positive cells from C using one-way ANOVA with Bonferroni’s post hoc test (n = 12-15, F (3, 52) = 57.31, p < 0.0001). Comparisons: DMSO vs. cortisol: ****p < 0.0001; DMSO vs. CORT125281 + cortisol: ****p < 0.0001; DMSO vs. CORT125281: p > 0.9999; cortisol vs. CORT125281 + cortisol: p > 0.9999, n.s., not significant. **(E)** Representative images illustrating the effects of cortisol on the morphology of Tuj1-positive neurons. **(F)** Statistics for the number of neurites from E using one-way ANOVA with Bonferroni’s post hoc test (n = 31-50, F (3, 150) = 7.777, p < 0.0001). Comparisons: DMSO vs. cortisol: ***p = 0.0009; DMSO vs. CORT125281 + cortisol: ***p = 0.0003; DMSO vs. CORT125281: p = 0.3250; cortisol vs. CORT125281 + cortisol: p > 0.9999, n.s., not significant. **(G)** Statistics for the maximum length of neurites from panel E, using one-way ANOVA with Bonferroni’s post hoc test (n = 31-62, F (3, 176) = 16.41, p < 0.0001). Comparisons: DMSO vs. cortisol: ****p < 0.0001; DMSO vs. CORT125281 + cortisol: ***p = 0.0005; DMSO vs. CORT125281: p = 0.8868; cortisol vs. CORT125281 + cortisol: p = 0.0840, n.s., not significant. **(H)** Statistics for the average length of neurites from E using one-way ANOVA with Bonferroni’s post hoc test (n = 13-16, F (3, 53) = 45.44, p < 0.0001). Comparisons: DMSO vs. cortisol: ****p < 0.0001; DMSO vs. CORT125281 + cortisol: *p = 0.0338; DMSO vs. CORT125281: p > 0.9999, n.s. = not significant; cortisol vs. CORT125281 + cortisol: ****p < 0.0001. Scale bars: 100 μm (A, C), 50 μm (E).

### In vivo blockade of glucocorticoid receptors promotes NSPCs proliferation and recovery post-SCI

These data suggest that elevated glucocorticoid levels following SCI suppress endogenous NSPC proliferation and differentiation, thereby hindering recovery. To investigate whether inhibition of glucocorticoid receptors enhances recovery, mice with SCI were orally treated with CORT125281 (30 mg/kg) or vehicle for three days (administered on days zero, one, and two post-SCI, Fig. 4A). CORT125281 treatment significantly increased the number of NSPCs at the injury site (Fig. 4B and C).

**Fig. 4.**
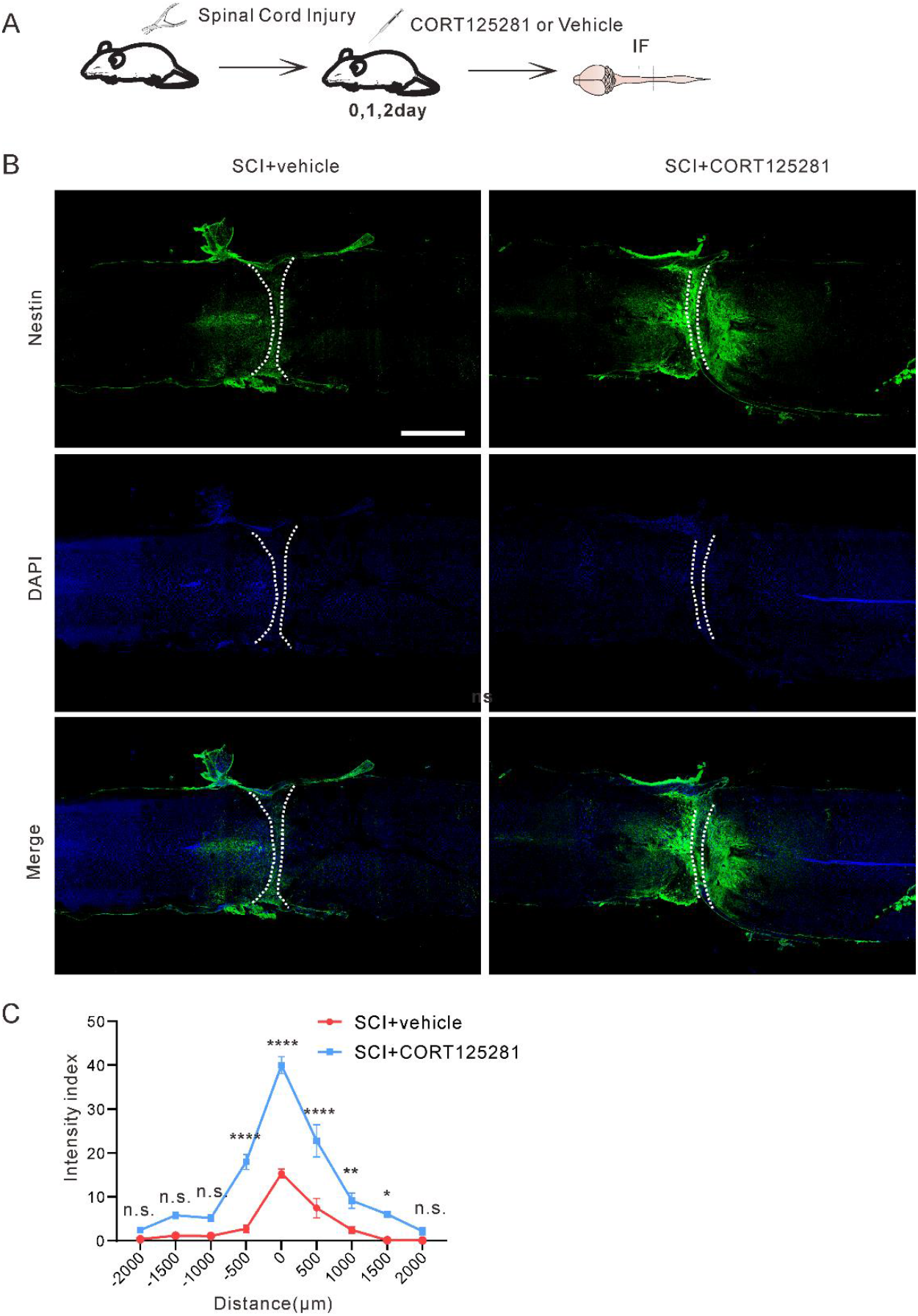
In vivo blockade of glucocorticoid receptors promotes NSPC proliferation post-SCI. **(A)** Schematic diagram illustrating the mouse spinal cord injury model, oral administration of the glucocorticoid receptor inhibitor, and spinal cord tissue collection. **(B)** Representative images showing nestin protein expression levels at the SCI site in mice. Scale bar: 1 mm. **(C)** Quantification of the density of nestin immunoreactive intensity (normalized to the density distal to the lesion site) from the lesion site, proximal to distal, in the spinal cord, two weeks post-injury. Data were analyzed using two-way ANOVA followed by Bonferroni’s post hoc test (n = 9 animals per group, F (8, 144) = 83.80, p < 0.0001). ****p < 0.0001, **p = 0.0055, *p = 0.0214, n.s., not significant.

Moreover, CORT125281 treatment led to marked improvements in motor function in the traumatic SCI mouse model. Mice receiving a placebo exhibited initial recovery of hip joint movement during the first two weeks post-surgery, with occasional foot-planting by the third week, but little further improvement in motor function thereafter. In contrast, mice treated with CORT125281 exhibited faster early recovery of hindlimb function. By the third week, they could consistently stand on their feet and showed continuous improvement over the following month. By weeks six to seven, some toe function had returned, and by week 9, some mice were able to walk with stable body movement (video 1 and 2, Fig. 5A). The BMS ^16^ was used to assess recovery after SCI. The BMS score of CORT125281-treated mice was significantly higher than that of the vehicle control mice (Fig. 5B). Additionally, in the grid-walking test, mice treated with the glucocorticoid receptor antagonist were able to place their hind paws correctly on the grid lines, exhibiting coordinated movement between forelimbs and hindlimbs. In contrast, placebo-treated mice showed minimal hindlimb movement and relied on their forelimbs to drag their bodies forward (Fig. 5C and D).

**Fig. 5.**
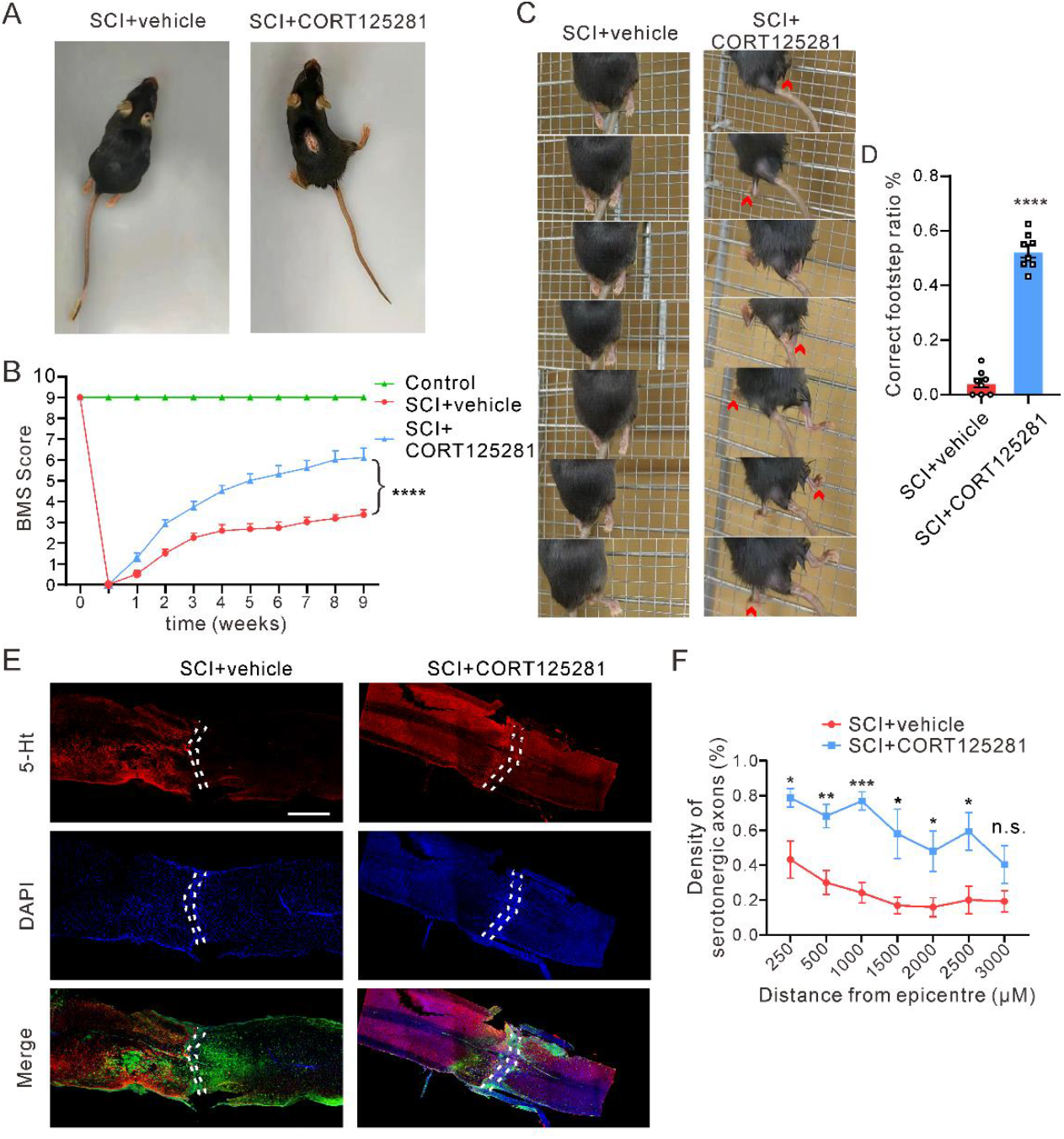
Oral administration of CORT125281 enhances motor function recovery in mice with SCI. **(A)** Representative videos showing the effect of vehicle control (video 1) or CORT125281 (video 2) on the recovery of motor function in mice, nine weeks post-SCI. **(B)** BMS score chart comparing motor function recovery in mice receiving CORT125281 or vehicle administration. Statistical analysis was performed using two-way ANOVA followed by Bonferroni’s post hoc test (Control: n = 10; SCI + vehicle: n = 11; SCI + CORT125281: n = 12), F (10, 221) = 175.8, p < 0.0001. **(C)** Grid climbing test results demonstrating the recovery of hindlimb motor function in the different groups of mice. **(D)** Statistics for the correct footstep ratio (number of correct hindfoot steps / number of correct forefoot steps) from panel C, using a two-tailed unpaired t-test (control: n = 8; CORT125281: n = 8), t (14) = 19.90, ****p < 0.0001. **(E)** Representative images of spinal sections stained with anti-serotonin (anti-5-HT) antibody, taken nine weeks after SCI, showing serotonergic axons. Dashed lines indicate the lesion site. Scale bar:1 mm. **(F)** Quantification of the density of serotonergic axons (normalized to the density proximal to the lesion site) in the spinal cord distal to the lesion site at nine weeks after injury. Statistical analysis was conducted using a two-tailed unpaired t-test (control: n = 6; CORT125281: n = 4), 250: t (8) = 2.537, *p = 0.0349; 500: t (8) = 3.762, **p = 0.0055; 1000: t (8) = 6.309, ***p = 0.0002; 1500: t (8) = 3.221, *p = 0.0122; 2000: t (8) = 2.803, *p = 0.0231; 2500: t (8) = 3.008, *p = 0.0169; 3000: t (8) = 1.863, p = 0.0995, n.s., not significant.

After nine weeks, we evaluated axon growth across the lesion site. Numerous serotonergic axons were observed in the spinal cord distal to the lesion in CORT125281-treated mice, in contrast to the vehicle-treated mice, where little or no axon regrowth was observed (Fig. 5E and F).

### Glucocorticoids induce NSPC arrest in the G0/G1 phase via glucocorticoid receptor-mediated regulation of cell cycle-related genes

Finally, we aimed to elucidate the molecular mechanisms underlying glucocorticoid-induced inhibition of adult NSPCs. ERK and PI3K/Akt signaling pathways play critical roles in neurogenesis ^17,18^. Therefore, we examined the effects of glucocorticoids on the activity of ERK and Akt in adult NSPCs. Treatment with 1 μM cortisol did not alter the activity of either ERK or Akt in adult NSPCs (Fig. 6A and B).

**Fig. 6.**
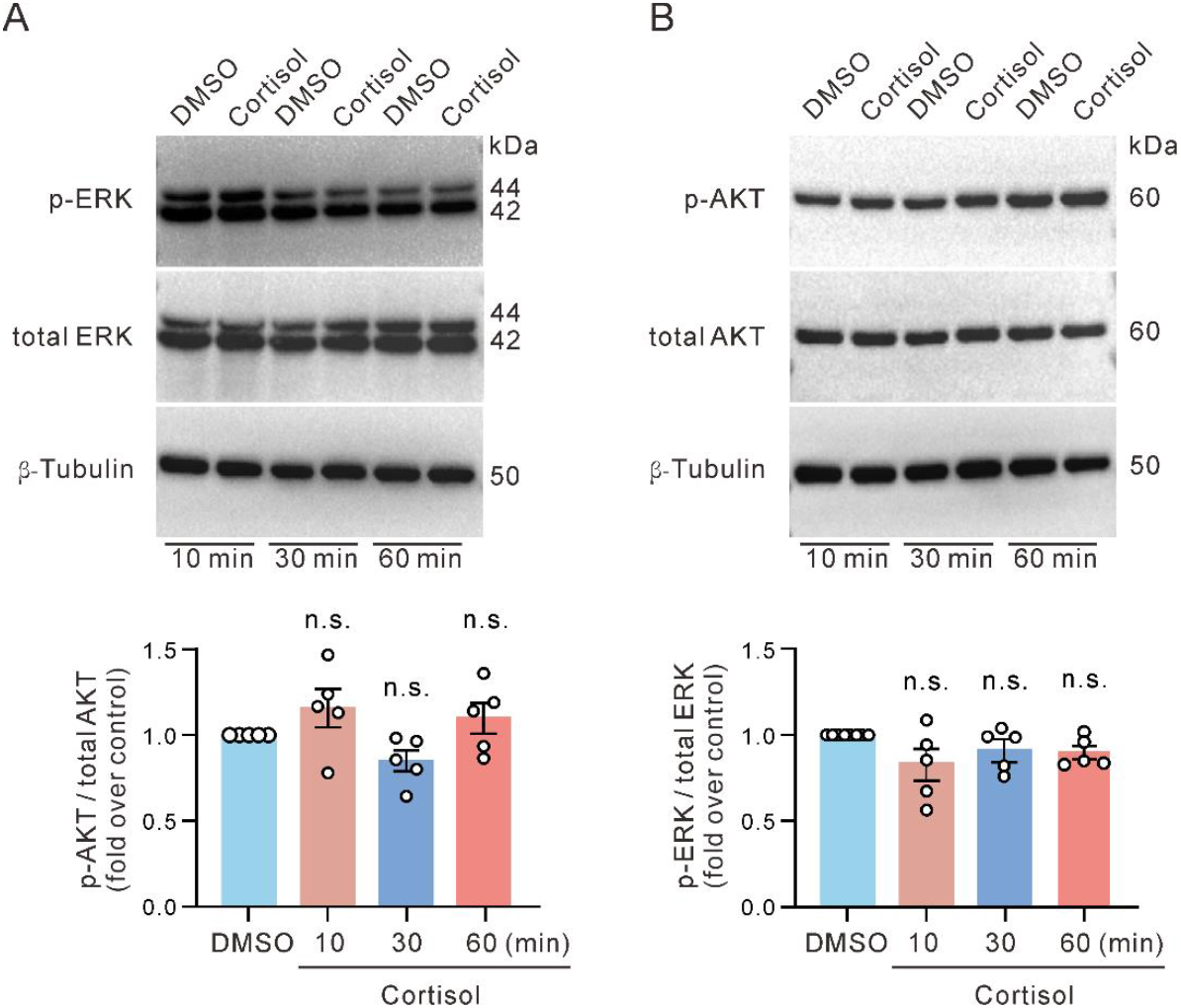
Cortisol does not alter the activity of ERK and AKT in adult NSPCs. **(A)** Top, Representative Western blot images show the effects of cortisol on phosphorylated ERK and total ERK in adult mouse NSPCs. β-Tubulin is used as a protein loading control. Bottom, Statistics from five independent experiments using one-way ANOVA with Bonferroni’s post hoc test. (n = 5, F (3, 16) = 1.839, p = 0.1808). 10 min: p = 0.1104; 30 min: p = 0.8490; 60 min: p = 0.3905, n.s., not significant. **(B)** Top, Representative Western blot images show the effects of cortisol on phosphorylated AKT and total AKT in adult mouse NSPCs. β-Tubulin is used as a protein loading control. Bottom, Statistics from five independent experiments using one-way ANOVA with Bonferroni’s post hoc test. (n = 5, F (3, 16) = 2.992, p = 0.0620). 10 min: p = 0.5157; 30 min: p = 0.5696; 60 min: p > 0.9999, n.s., not significant.

To further explore the underlying mechanism of glucocorticoid-induced inhibition of adult NSPCs, RNA sequencing was performed on cultured adult spinal cord NSPCs treated with 1 μM cortisol for three days. Comparative analysis of mRNA expression profiles between NSPCs treated with DMSO and those treated with cortisol revealed a total of 222,098 transcripts. Of these, 366 mRNAs were significantly altered by at least two-fold in cortisol-treated cells. Specifically, 232 mRNAs were upregulated, while 134 were downregulated relative to DMSO controls (Fig. 7A-C). KEGG pathway classification analysis categorized the differentially expressed genes involved in various cellular processes (Fig. 7D). Among these, genes related to cell death, growth, and the nervous system were of particular interest in this study. Further KEGG pathway enrichment analysis revealed that the p53 signaling pathway was implicated in glucocorticoid-mediated effects on NSPCs (Fig. 7E). To validate the RNA sequencing results, we confirmed the top five upregulated and downregulated mRNAs in cortisol-treated NSPCs using qRT-PCR (Fig. 7F and G). The qRT-PCR results confirmed sequencing findings.

**Fig. 7.**
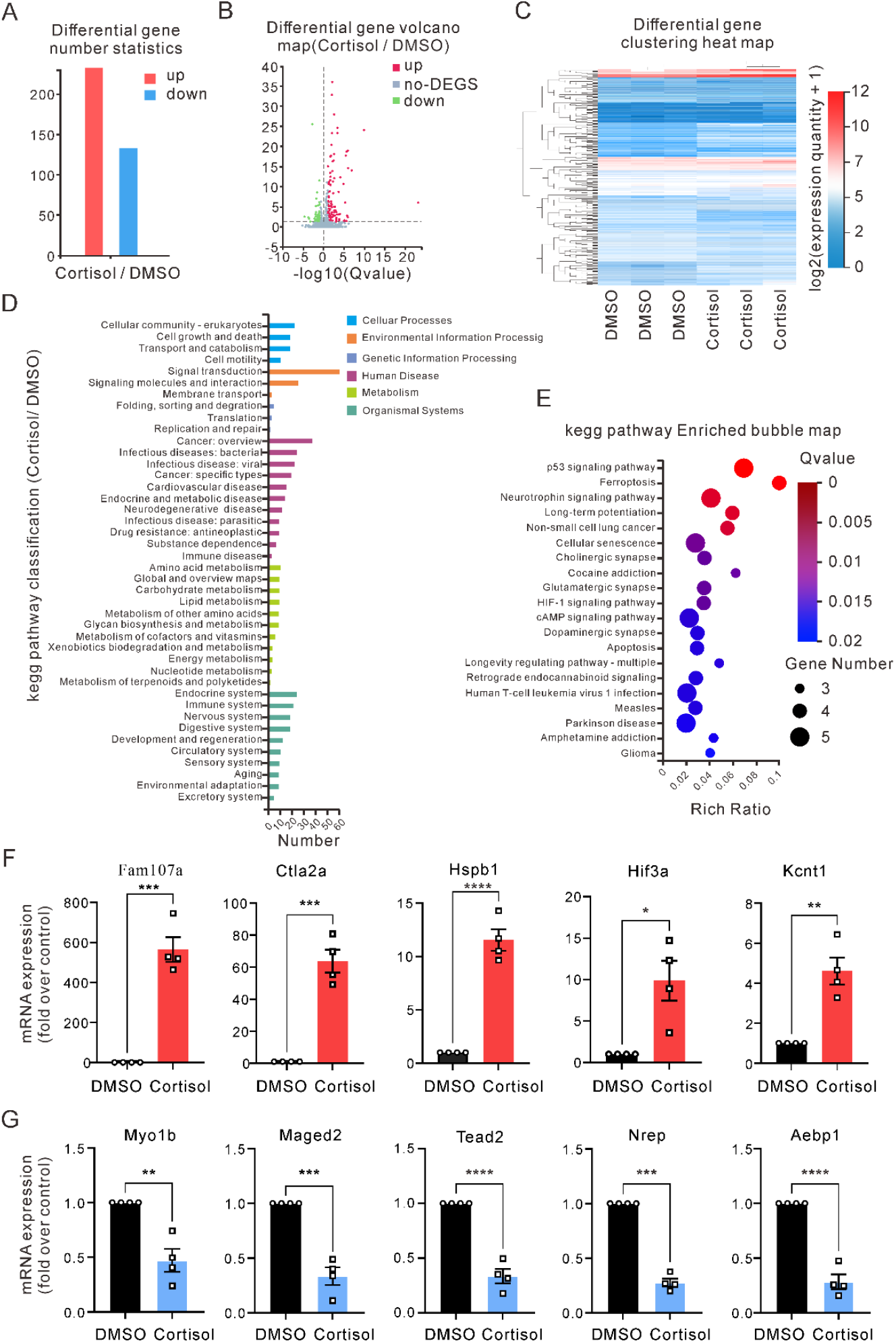
RNA sequencing analysis of gene expression changes in NSPCs following cortisol treatment. **(A)** Statistical chart showing differentially expressed genes with more than a twofold change in expression. **(B)** Volcano plot of differentially expressed genes. **(C)** Heatmap of clustered differentially expressed genes. **(D)** KEGG pathway classification of all differentially expressed genes. **(E)** KEGG pathway enrichment analysis of genes of interest, with the p53 signaling pathway identified as the most relevant. **(F, G)** qRT-PCR validation of mRNA sequencing results. Data were analyzed using a two-tailed unpaired t-test (n = 4). *Fam107a*: t (6) = 9.080, ***p = 0.0001; *Ctla2a*: t (6) = 8.801, ***p = 0.0001; *Hspb1*: t (6) = 10.50, ****p < 0.0001; *Hif3a*: t (6) = 3.694, *p = 0.0102; *Kcnt1*: t (6) = 5.369, **p = 0.0017; *Aebp1*: t (6) = 10.62, ****p < 0.0001; *Tead2*: t (6) = 10.18, ****p < 0.0001; *Maged2*: t (6) = 8.243, ***p = 0.0002; *Nrep*: t (6) = 19.89, ****p < 0.0001; *Myo1b*: t (6) = 5.040, **p = 0.0024.

Given the significant role of p53 signaling in cell cycle regulation, we investigated the effects of cortisol on the NSPC cell cycle. Flow cytometry results revealed that cortisol induced cell cycle arrest at the G1/G0 phase (Fig. 8A and B). The progression of cells through the cell cycle is regulated by a family of protein kinases known as cyclin-dependent kinases. In mammals, the cyclin-dependent kinase inhibitors are divided into two families: the Ink family, which includes p16, p15, p18, and p19, and the Cip/Kip family, which comprises p21, p27, and p57 ^19^. Cortisol significantly increased the expression of p15, p16, p18, and p27, an effect that could be reversed by the glucocorticoid receptor inhibitor, CORT125281 (Fig. 8C). In contrast, cortisol had no effect on the expression of p19, p21, or p57 (Fig. 8C).

**Fig. 8.**
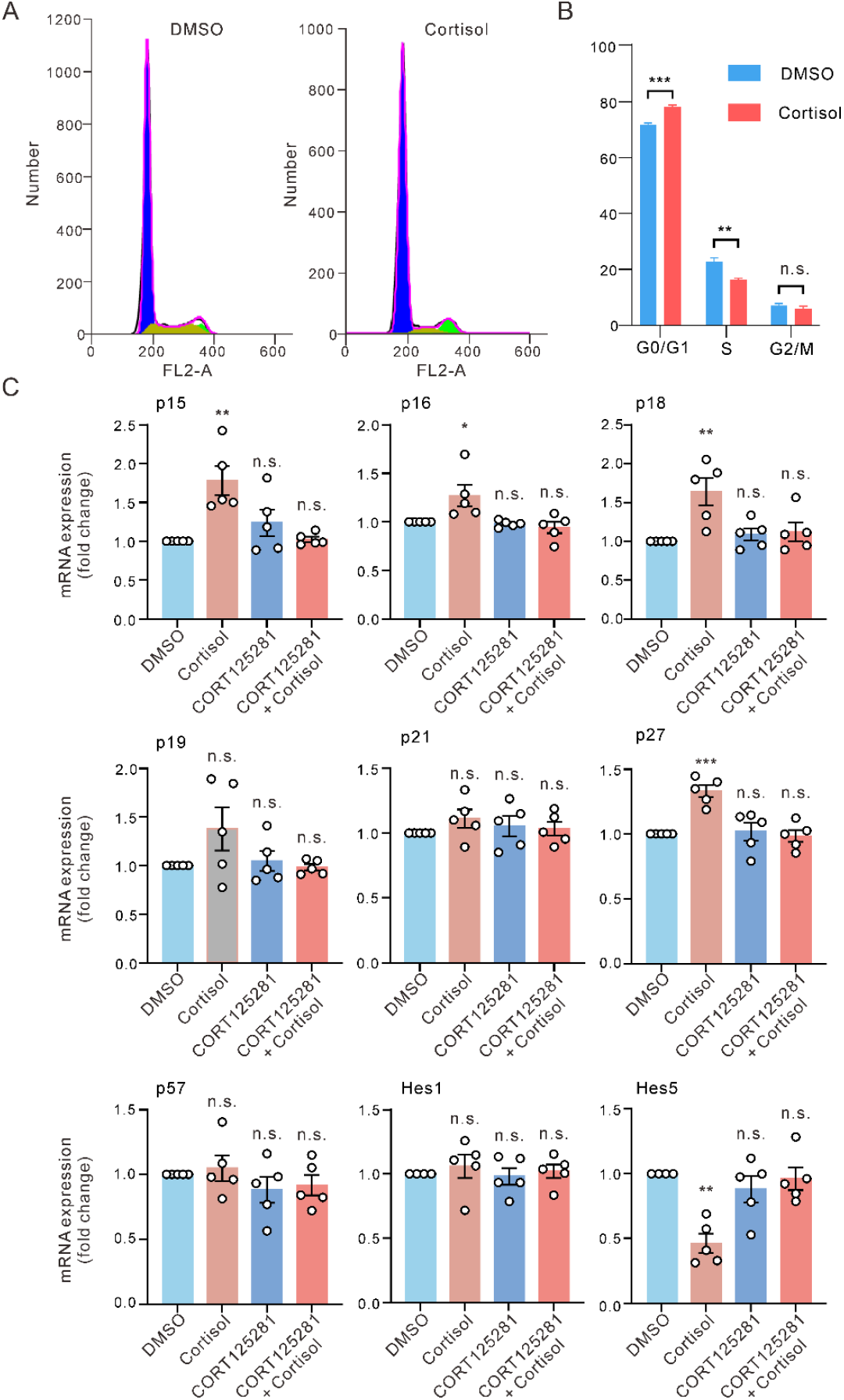
Glucocorticoids induce NSPC arrest in the G0/G1 phase via glucocorticoid receptor-mediated regulation of cell cycle-related genes. **(A)** Representative flow cytometry histograms showing the distribution of cells in different phases of the cell cycle (G0/G1, S, and G2/M) after treatment with 1μM cortisol for 36 hours. FlowJo software was used for histogram analysis. **(B)** Statistical analysis revealed significant differences in cell cycle phases: G0/G1 phase (t (4) = 13.03, ***p = 0.0002), S phase (t (4) = 6.874, **P = 0.0023), and G2/M phase (t (4) = 1.016, p = 0.3669, n.s., not significant). Two-tailed unpaired t-test, n = 3. **(C)** The effect of 1 μM cortisol on mRNA expression of cell cycle-related genes (p15, p16, p18, p19, p21, p27, p57) and neural stem cell maintenance genes (Hes1, Hes5). Data were analyzed using one-way ANOVA with Bonferroni’s post hoc test (n = 5). *p15*: (F (3, 16) = 8.012, p = 0.0017). DMSO vs. cortisol: **p = 0.0016; DMSO vs. CORT125281 + cortisol: p > 0.9999; DMSO vs. CORT125281: p = 0.6345. *p16*: (F (3, 16) = 5.586, p = 0.0081). DMSO vs. cortisol: *p = 0.0259; DMSO vs. CORT125281 + cortisol: p > 0.9999; DMSO vs. CORT125281: p > 0.9999. *p18*: (F (3, 16) = 6.578, p = 0.0042). DMSO vs. cortisol: **p = 0.0031; DMSO vs. CORT125281 + cortisol: p > 0.9999; DMSO vs. CORT125281: p = 0.6345. *p19*: (F (3, 16) = 2.245, p = 0.1225). DMSO vs. cortisol: p = 0.1417; DMSO vs. CORT125281 + cortisol: p > 0.9999; DMSO vs. CORT125281: p > 0.9999. *p21*: (F (3, 16) = 0.6257, p = 0.6077). DMSO vs. cortisol: p = 0.6029; DMSO vs. CORT125281 + cortisol: p > 0.9999; DMSO vs. CORT125281: p > 0.9999. *p27*: (F (3, 16) = 12.21, p = 0.0002). DMSO vs. cortisol: p = 0.0005; DMSO vs. CORT125281 + cortisol: p > 0.9999; DMSO vs. CORT125281: p > 0.9999. *p57*: (F (3, 16) = 0.8859, p = 0.4694). DMSO vs. Cortisol: p > 0.9999; DMSO vs. CORT125281 + cortisol: p = 0.9497; DMSO vs. CORT125281: p > 0.9999. *Hes1*: (F (3, 16) = 0.2667, p = 0.8483). DMSO vs. cortisol: p > 0.9999; DMSO vs. CORT125281 + cortisol: p > 0.9999; DMSO vs. CORT125281: p > 0.9999. *Hes5*: (F (3, 16) = 9.420, p = 0.0010). DMSO vs. cortisol: **p = 0.0012; DMSO vs. CORT125281 + cortisol: p = 0.9784; DMSO vs. CORT125281: p > 0.9999; n.s., not significant.

Hes1 and Hes5 genes play essential roles in the maintenance and proliferation of neural stem cells^20-22^. We also assessed the effect of cortisol on these genes. Real-time PCR results showed that 1 μM cortisol significantly decreased the mRNA expression of Hes5, while it did not alter the expression of Hes1 (Fig. 8C).

## Discussion

Glucocorticoid levels significantly increase following spinal cord injury (SCI), but their effects on adult spinal cord neural stem/progenitor cells (NSPCs) remains poorly understood. In this study, we demonstrate that glucocorticoids suppress the proliferation of adult spinal cord NSPCs through glucocorticoid receptors. Glucocorticoids induce NSPC arrest in the G0/G1 phase via glucocorticoid receptor-mediated regulation of cell cycle-related genes. Moreover, treatment with the glucocorticoid receptor inhibitor CORT125281 significantly enhanced motor function in a traumatic SCI mouse model. These results suggest that targeting glucocorticoid receptors with specific inhibitors may offer a novel therapeutic strategy to promote recovery after SCI.

Endogenous glucocorticoids bind to two receptors: the mineralocorticoid receptor, which has a high affinity, and the glucocorticoid receptor, which is activated at higher glucocorticoid levels ^23^. Both receptors are expressed in embryonic NSPCs. Typically, glucocorticoids inhibit the proliferation of embryonic NSPCs via glucocorticoid receptors ^24^. Anacker et al. reported that a low concentration of cortisol (100 nM) enhanced the proliferation of human hippocampal progenitor cells via the mineralocorticoid receptor, whereas a high concentration of cortisol (100 μM) reduced proliferation by activating glucocorticoid receptors ^25^. In contrast, we found that adult spinal cord NSPCs express only the glucocorticoid receptor, and glucocorticoids inhibit their proliferation in a dose-dependent manner via this receptor.

Previous studies have shown that melatonin promotes the proliferation of adult spinal cord NSPCs through the PI3K/AKT signaling pathway ^26^. Similarly, Lu et al. reported that icariin sustains the proliferation of Aβ-treated hippocampal neural stem cells via the ERK/Akt signaling pathway ^27^. However, we found that glucocorticoids did not alter ERK or Akt signaling. RNA sequencing analysis further revealed that the p53 signaling pathway is involved in the glucocorticoid-mediated suppression of adult spinal cord NSPC proliferation.

p53 is known as a key regulator of cell growth and division, and its activation leads to G1 arrest ^28^. Previous studies have demonstrated that glucocorticoid receptors and p53 interact both physically and functionally, with such interactions contributing to various physiological and pathological outcomes ^29,30^. Dexamethasone has been shown to induce G0/G1 arrest and increase p21 expression in MC3T3-E1 cells through aberrant glucocorticoid receptor activation, leading to subsequent p53 activation ^31^. In addition, dexamethasone induces G1-phase arrest and increases p27 expression in hippocampal progenitor cells ^32^. Similarly, dexamethasone upregulates cell-cycle regulatory genes such as p16 and p21 in rat embryonic neural stem cells^10^. In this study, we observed that glucocorticoids induce G1-phase arrest and increase the expression of cell-cycle regulators p15, p16, p18, and p27 in adult spinal cord NSPCs. The effects of glucocorticoids on the differentiation of adult spinal cord NSPCs are complex. Our findings show that glucocorticoid receptor antagonists can partially block glucocorticoid-induced inhibition of glial cell differentiation but have no effect on the suppression of neuronal differentiation. These results suggest that additional mechanisms beyond the classical glucocorticoid receptor pathway may mediate the effects of glucocorticoids on the neuronal differentiation of adult spinal cord NSPCs.

Changes in the microenvironment following SCI impact the balance between inhibitory and growth-promoting factors^8^. For example, after SCI, the expression of myelin-associated inhibitors, such as Nogo-A, increases, further impeding axonal regeneration^33^. In contrast, growth-promoting molecules like neurotrophin-3 may be downregulated, limiting their availability to injured neurons ^34^. Researchers are exploring strategies such as targeting inflammatory pathways, degrading inhibitory molecules, and enhancing the availability of growth-promoting factors to create a more favorable microenvironment for recovery ^35^. High-dose methylprednisolone (a synthetic glucocorticoid) is commonly administered in clinical settings for the treatment of acute SCI. However, an increasing body of evidence suggests that methylprednisolone fails to improve motor function or neurological outcomes, while also causing notable adverse effects ^36-38^. Furthermore, studies have indicated that clinically relevant doses of methylprednisolone may inhibit the proliferation of endogenous spinal cord NSPCs following SCI ^39,40^. Recent studies have shown that glucocorticoid levels are significantly elevated in both murine and human SCI models^9^. In our study, we found that glucocorticoids suppress the proliferation of endogenous adult spinal cord NSPCs through glucocorticoid receptors, impairing SCI recovery.

In conclusion, our study provides the first evidence that glucocorticoids inhibit the proliferation and differentiation of endogenous adult spinal cord NSPCs, and that blocking glucocorticoid receptors enhances motor function in a traumatic SCI model in mice. This study offers a potential therapeutic strategy with the promise of functional recovery after SCI.

## Materials and Methods

### Ethics statement

All protocols performed in studies involving animal were approved by the Committee on the Ethics of Animal Experiments of Fudan University and were in strict accordance with the NIH Guidelines for the Care and Use of Laboratory Animals.

### Reagents and chemicals

Dulbecco’s Modified Eagle’s Medium (DMEM)/F12, B27, N2, and fetal bovine serum (FBS) were obtained from Gibco (Grand Island, NY, USA). Accutase, protease inhibitor cocktail, and phosphatase inhibitor cocktail 2 were purchased from Sigma-Aldrich (Munich, Germany). Basic fibroblast growth factor (bFGF) and epidermal growth factor (EGF) were sourced from Peprotech (Rocky Hill, NJ, USA). Corticosterone and cortisol were purchased from Sigma (St. Louis, MO, USA). CORT125281 was obtained from MCE (Brooklyn, NY, USA).

### Primary culture of adult mouse spinal cord stem/progenitor cells

Six-week-old female C57BL/6 mice were purchased from Slack (Shanghai, China). Adult spinal cord NSPCs were isolated as previously described ^26^. The cells were cultured in proliferation medium consisting of DMEM/F12 supplemented with 1% B27, 1% N2, 1% Penicillin-Streptomycin, 10 ng/mL EGF, and 20 ng/mL bFGF. NSPCs at passages one to four were used for the experiments. For differentiation assays, the cells were cultured in differentiation medium containing DMEM/F12 supplemented with 1% B27, 1% N2, 1% Penicillin-Streptomycin, and 1% FBS.

### Neurosphere assay

Adult mouse spinal cord NSPCs were seeded in 96-well plates at a density of 5,000 cells per well and cultured in proliferation medium with or without various drug treatments for seven days. Neurospheres were considered to be of standard size when measuring 70 µm in diameter. The number of neurospheres was quantified using ImageJ software (National Institutes of Health, MD, USA).

### Cell viability assay

Cell viability was assessed using the Cell Counting Kit-8 (CCK-8) following the manufacturer’s instructions (Beyotime, Shanghai, China). Briefly, neurospheres were dissociated into single cells and cultured in 96-well plates with or without cortisol for five days. Subsequently, cells were incubated with CCK-8 solution for four hours at 37°C. The optical density (OD) was measured at 450 nm to determine cell viability.

### 5-Ethynyl-2-Dexoyuridine (EdU) incorporation assay

Spinal cord NSPCs were cultured in proliferation medium with or without cortisol for 24 hours, then dissociated into single cells and plated onto poly-L-ornithine-coated glass slides. After an eight-hour incubation in proliferation medium containing EdU, the cells were fixed with 4% paraformaldehyde (PFA) at room temperature for 30 minutes. The PFA was then removed, and the cells were incubated with a glycine solution for six minutes. After two washes with PBS, the cells were permeabilized with 0.5% Triton X-100 in PBS for ten minutes. Subsequently, the cells were incubated with the staining mix for 30 minutes in the dark at room temperature, followed by three washes with PBS. Cell nuclei were counterstained with DAPI. The ratio of EdU-positive cells to DAPI-positive cells was calculated under fluorescence microscopy. For in vivo analysis, EdU (50 mg/kg) was administered daily from day three to day five post-SCI. On day five following injury, the animals were anesthetized as previously described and perfused with 4% PFA. EdU incorporation was detected using the BeyoClick™ EdU-647 Cell Proliferation Kit (Beyotime, Shanghai, China).

### Quantitative real-time polymerase chain reaction (qRT-PCR)

Spinal cord NSPCs were cultured in proliferation medium with or without cortisol for five days. Total RNA was extracted, and qRT-PCR was performed as previously described ^41^. Briefly, qRT-PCR was conducted in 20 μL reactions containing 2 μL of cDNA template, 0.4 μmol/L of each primer pair, and SYBR Green PCR Master Mix (TIANGEN, Beijing, China). The thermal cycling conditions were as follows: 94°C for 10 minutes, followed by 40 cycles of 94°C for 30 seconds, 55°C for 30 seconds, and 72°C for 30 seconds, with a final extension at 72°C for eight minutes. Results were normalized to β-actin mRNA. Data were analyzed using the 2^-ΔΔCt^ method and are expressed as fold change relative to the control. The primer sequences are listed in Supplementary Table 1.

### Western blot

NSPCs were cultured in proliferation medium and treated with cortisol for 10, 30, and 60 minutes. Following treatment, total cellular protein was extracted using ice-cold RIPA buffer containing protease and phosphatase inhibitors to preserve protein integrity. The protein samples were resolved using 10% SDS PAGE and transferred to polyvinylidene fluoride membranes. The membranes were immunoblotted with primary antibodies against phospho-ERK (p-ERK, 1:1000, #4370, Cell Signaling Technology), phospho-AKT (p-AKT, 1:1000, #4060, Cell Signaling Technology), total ERK (1:1000, #4695, Cell Signaling Technology), total AKT (1:1000, #9272, Cell Signaling Technology), and β-tubulin (1:1000, #AF2835, Beyotime) overnight at 4°C. After washing with TBST three times, the membranes were incubated with HRP-conjugated secondary antibody (goat anti-mouse, 1:1000, #P0946, Beyotime; goat anti-rabbit, 1:1000, #P0948, Beyotime) for one hour at room temperature. The blots were developed using enhanced chemiluminescence reagents and imaged using the ChemiDoc XRS+ imaging system (Bio-Rad, Hercules, CA, USA) with the manufacturer’s software. Band intensities were quantified using ImageJ software. Phosphorylation levels of ERK and AKT were normalized to their corresponding total protein levels.

### Immunofluorescence

Neurospheres or single cells seeded on poly-L-ornithine-coated slides were fixed with 4% PFA for 30 minutes at room temperature, followed by three washes with PBS. Cells were then incubated with a blocking solution containing 10% horse serum and 0.5% Triton X-100 in PBS for one hour at room temperature. After three PBS washes, cells were incubated overnight at 4°C with primary antibodies: anti-GR (1:1000, #3660, Cell Signaling Technology), anti-MR (1:500, ab64457, Abcam), anti-GFAP (1:1000, ab7260, Abcam), anti-Tuj1 (1:1000, ab18207, Abcam), anti-nestin (1:1000, ab6142, Abcam), anti-5-HT (1:500, #20079, IMMUNOSTAR). Following three washes with PBST (0.025% Triton X-100 in PBS), cells were incubated with secondary antibodies (Alexa Fluor® 488-conjugated donkey anti-mouse IgG, 1:500, 715-545- 150, Jackson; Cy5-conjugated donkey anti-goat IgG, 1:500, 705-175-147, Jackson; Alexa Fluor® 647- conjugated goat anti-rabbit, 1:500, #A0468, beyotime) in 1% horse serum and 0.3% Triton X-100 in PBS overnight at 4°C. After washing with PBS three times, cells were counterstained with DAPI. The chest region of the spinal cord was used to analyze the organization and distribution of NSCs in response to SCI. Mice were anesthetized with pentobarbital and subsequently transcardially perfused with ice-cold 4% PFA in PBS. The spinal cord was then carefully excised and post-fixed in the same fixative at 4°C for four to eight hours. After fixation, the tissue was equilibrated in 30% (w/v) sucrose in PBS overnight at 4°C, embedded in optimal cutting temperature compound, and frozen for sectioning. Immunofluorescence images were captured using a Nikon confocal microscope.

### Flow cytometry analysis

Cells were collected at specified time points and fixed. The cell pellet was resuspended and incubated with RNase A before being stained with propidium iodide. Cell cycle distribution was then evaluated using flow cytometry.

### SCI model

To model clinical compression injury of the spinal cord, a crush-injured SCI model was established as previously described ^42^. Adult female C57BL/6J mice (16–20 g, 6 weeks of age) were selected to ensure uniform spinal cord size. To minimize lesion variability due to differences in surgical experience, all SCI procedures were performed by the same surgeon. Briefly, animals were anesthetized with 2% pentobarbital (30 mg/kg) and a ∼2 cm incision was made along the midline of the back. After performing a laminectomy at the T8 level, the spinal cord was laterally compressed for 15 seconds using a pair of specialized forceps with a 0.4 mm spacer, producing a precise moderate crush injury.

### Assessment of locomotor performance

Hindlimb function in mice was assessed weekly post-surgery using the Basso Mouse Scale (BMS) open-field locomotor test ^16^ and the inclined-grid climbing test ^43^. The BMS test quantitatively evaluates voluntary movement and the ability to support body weight. The inclined-grid climbing test assesses the percentage of mice that regained the ability to place their hindlimbs correctly and recover proprioceptive responses.

### RNA sequencing

NSPCs were collected after treatment under different conditions. Total RNA was isolated using the RNeasy Kit (Qiagen, Germany) according to the manufacturer’s instructions. The extracted RNA was used to construct RNA sequencing libraries, which underwent quality control analysis before sequencing on the DNBSEQ platform. Clean reads were mapped to the reference genome using HISAT2 (v2.0.4). Bowtie2 (v2.2.5) was employed to align the clean reads to the reference coding gene set, and gene expression levels were calculated using RSEM (v1.2.12). Differential gene expression analysis was performed using DESeq2 (v1.4.5). GO and KEGG enrichment analyses were conducted using Phyper, based on the Hypergeometric test, with statistical significance corrected by the Bonferroni method at a stringent threshold (Q value ≤ 0.05). The raw sequencing data included in this study has been deposited in the SRA database (https://www.ncbi.nlm.nih.gov/sra) under the accession code PRJNA1208143.

### Statistics

All statistical analyses were performed using GraphPad Prism (v9.4, GraphPad Software Inc, USA). The normality of the data was assessed using the Shapiro–Wilk test. The two tailed paired or unpaired *t* test was used to compare two groups, and one-way or two-way ANOVA with Bonferroni’s post hoc test was employed for comparisons involving multiple groups. Data are presented as mean ± SEM, with “n*”* representing the number of independent experiments or animals. A p-value of < 0.05 was considered statistically significant.

## CRediT authorship contribution statement

X.Z.: Investigation, Formal analysis, Writing – review & editing. S.Z., X.H., S.T., Y. C.: Investigation. C.H.: Conceptualization, Supervision, Writing – original draft, Writing –review & editing.

## Data availability

All data is available upon request from the corresponding author. The raw sequencing data included in this study has been deposited in the SRA database (https://www.ncbi.nlm.nih.gov/sra) under the accession code PRJNA1208143.

## Acknowledgements

This work was supported by the National Key Research and Development Program of China (2022YFC3602702), the Science and Technology Innovation 2030 Brain Science and Brain-Inspired Intelligence Project (2021ZD0201301), and the Natural Science Foundation of Shanghai (23ZR1425900)

