## Supplemental Table 1 for "Inhibiting glucocorticoid receptors enhances adult spinal cord neural stem cell activity and improves outcomes in spinal cord injury"

**Supplemental table 1: The primer sequences for the qRT-PCR.**

| Gene | Forward primer | Reverse primer |
|---|---|---|
| Fam107a | 5'- CAGACCAGAGTACAGAGAGTGG | 5'- GTGGTTCATAAGCAGCTCACG |
| Ctla2a | 5'- CTCCACCCCCTGATCCAAGT | 5'- ACACGAGTCTTCTGTGTCTTTCT |
| Hspb1 | 5'- ATCCCCTGAGGGCACACTTA | 5'- GGAATGGTGATCTCCGCTGAC |
| Hif3a | 5'- GAAGTTCACATACTGCGACGA | 5'- GTCCAAAGCGTGGATGTATTCAT |
| Kcnt1 | 5'- TGAGGTGACACCCTCGGAC | 5'- GGTGGCATAAAGTCGGGGAG |
| Myo1b | 5'- GGAGGGAGTTGAAACGCTTGA | 5'- CAGTAAGCCCAAATGACAGCAA |
| Maged2 | 5'- TCTGACACAAGCGAGAGTGG | 5'- TCAACAGGTCCTGCATCACG |
| Tead2 | 5'- GAAGACGAGAACGCGAAAGC | 5'- GATGAGCTGTGCCGAAGACA |
| Nrep | 5'- GTCAGCCAAGAACCGTTTGC | 5'- TTCACTTCCTTAGGCACGGG |
| Aebp1 | 5'- TTGGAAACGCTGGATCGGTTA | 5'- CTTGACCTTGCCAGGCATTT |
| p15 | 5'- CCCTGCCACCCTTACCAGA | 5'-GCAGATACCTCGCAATGTCAC |
| p16 | 5'-CGCAGGTTCTTGGTCACTGT | 5'-TGTTCACGAAAGCCAGAGCG |
| p18 | 5'- CCTTGGGGGAACGAGTTGG | 5'-AAATTGGGATTAGCACCTCTGAG |
| p19 | 5'-CTGGAAGAAGTCTGCGTCGG | 5'-GTCTTGCCAAAGCGGTTCAG |
| p21: | 5'- CCTGGTGATGTCCGA CCTG | 5'- CCATGAGCGCATCGCAATC |
| p27 | 5'-TCAAACGTGAGAGTGTCTAACG | 5'- CCGGGCCGAAGAGATTTCTG |
| p57: | 5'- CGAGGAGCAGGACGAGAATC | 5'- GAAGAAGTC GTTCGCATTGGC |
| hes1 | 5'- GAAGAGGCGAAGGGCAAGAA | 5'- GAGGTGCTTCACAGTCATTTCCA |
| hes5 | 5'- GACCGCATCAACAGCAGCAT | 5'- GGCGAAGGC TTTGCTGTGT |
| β-actin | 5′-GGACTTCGAGCAAGAGATGG | 5′ -AGCACTGTGTTGGCGTACAG |
